# Optimized replication of arrayed bacterial mutant libraries increase access to biological resources

**DOI:** 10.1101/2023.04.25.537918

**Authors:** Julia L. E. Willett, Aaron M. T. Barnes, Debra N. Brunson, Alexandre Lecomte, Ethan B. Robertson, Gary M. Dunny

## Abstract

Biological collections, including arrayed libraries of single transposon or deletion mutants, greatly accelerate the pace of bacterial genetics research. Despite the importance of these resources, few protocols exist for the replication and distribution of these materials. Here, we describe a protocol for creating multiple replicates of an arrayed bacterial Tn library consisting of approximately 6,800 mutants in 73 × 96-well plates. Our protocol provides multiple checkpoints to guard against contamination and minimize genetic drift caused by freeze/thaw cycles. This approach can also be scaled for arrayed culture collections of various sizes. Overall, this protocol is a valuable resource for other researchers considering the construction and distribution of arrayed culture collection resources for the benefit of the greater scientific community.

**Importance:** Arrayed mutant collections drive robust genetic screens, yet few protocols exist for replication of these resources and subsequent quality control. Increasing distribution of arrayed biological collections will increase accessibility to and use of these resources. Developing standardized techniques for replication of these resources is essential for ensuring their quality and usefulness to the scientific community.

## Introduction

Mutagenesis of a given organism followed by phenotypic selection or measurement of mutant fitness is a cornerstone of experimental microbial genetics. High-quality, publicly available collections of mutants, such as the *Escherichia coli* KEIO collection (1), *Bacillus subtilis* single-gene knockout libraries (2), and the *Staphylococcus aureus* USA300_FPR3757 transposon (Tn) mutant library (3) greatly accelerate the pace at which research can be done and enhance scientific rigor and reproducibility. Construction of such resources requires significant time, labor, and resources, and it is inefficient for multiple laboratories to generate redundant biological resources. The research community would benefit from increased generation, replication, and dissemination of resources like arrayed bacterial mutant libraries. Although robust protocols have been developed for manual or robotic arraying of colonies and mapping arrayed collections of mutants (4-8), few exist for the replication of arrayed culture collections (9-11). Here, our goal was to establish a protocol and collection of best practices to minimize contamination and genetic drift of arrayed bacterial culture collections while increasing accessibility to other researchers.

We previously described the generation and application of an arrayed library of Tn mutants in *Enterococcus faecalis* OG1RF consisting of ∼15,000 individual clones (12). From this library, two targeted SmarT (Sequence-defined *mariner* Transposon) libraries were generated (5). The first SmarT library consists of 6,829 Tn mutants arrayed across 73 96-well plates with insertions in approximately 70% of annotated genes and intergenic regions in OG1RF. The second SmarT library consists of 1,946 Tn insertions in poorly characterized genes and intergenic regions and was designed to facilitate genetic screens targeting uncharacterized regions of the genome (13). Both libraries are also available in pooled formats and have been used extensively to identify *E. faecalis* genes required for biofilm formation, metabolism, response to antibiotics, phage infection, vaginal colonization in a mouse model, and polymicrobial interactions involving *E. faecalis* (14-21).

In addition to these genetic screens, hundreds of individual Tn mutants have been distributed to domestic and international labs. We regularly receive requests for the entire Tn library, but it is not feasible to generate a new copy of the entire arrayed library for each individual request. Therefore, we sought to do a large-scale replication of the larger SmarT library (6,829 mutants) to increase accessibility of this resource by other labs, ensure quality control of the collection, and avoid genetic drift by decreasing the number of freeze/thaw cycles for the original library plates. Here, we present a protocol for efficient manual replication of arrayed library resources, including estimation of the time required (person hours). This protocol does not require access to robotic handling systems, making it feasible for researchers that do not have access to this specialized equipment. This protocol can be scaled to accommodate libraries of different sizes as well as different numbers of replicates. We also describe multiple quality control checks throughout the process and compare sequencing-based verification of pooled mutants with previously published results. Additionally, because of ongoing supply chain difficulties due to the COVID-19 pandemic, we consider multiple options for consumables required throughout as well as ergonomic considerations for technical staff.

## Results

### Abbreviated protocol for large-scale replication and quality control of arrayed transposon resources

We manually created 15 copies of the 73-plate SmarT library over a two-week period in April 2022. Required supplies and consumables are listed in **Table 1**. The approximate timeline for library replication of this scale is outlined in **Table 2** and **Table 3**. Additional protocol details can be found in **Supplementary File 1**. To avoid repeated freeze/thaw cycles of the original library plates, all copies were created at the same time. An overview of the process is shown in **Figure 1**. Frozen SmarT library stock plates were used to inoculate deep 96-well plates containing BHI. Cultures were grown overnight and manually inspected for contamination of known blanks or lack of growth. If plates had either contaminated wells or wells with no growth, the entire plate was discarded, and a new deep-well plate was inoculated. Overnight cultures were transferred from deep-well plates to pre-labeled 96-well plates containing glycerol to generate individual library sets and stored at -80 °C.

**Table 1.**
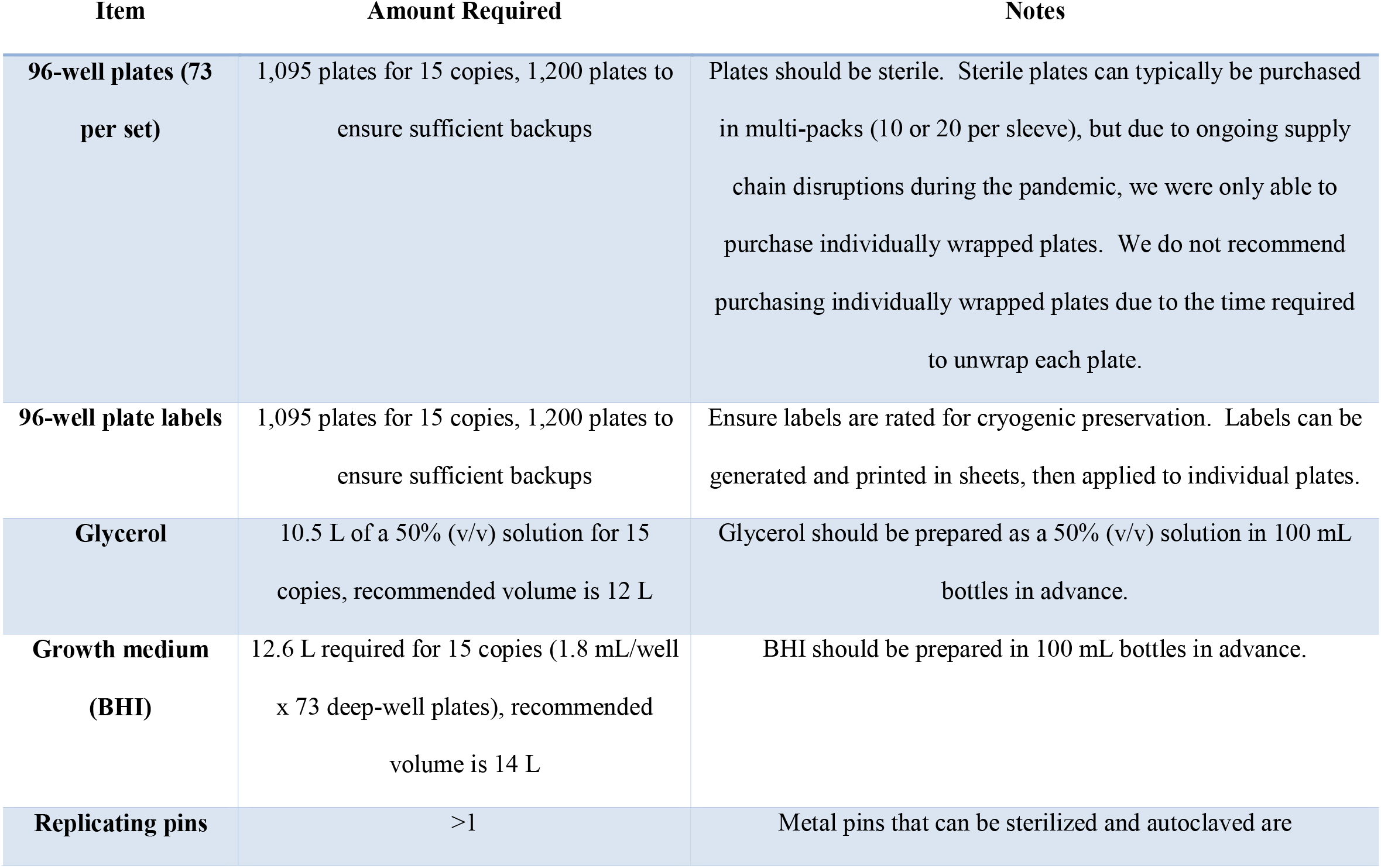
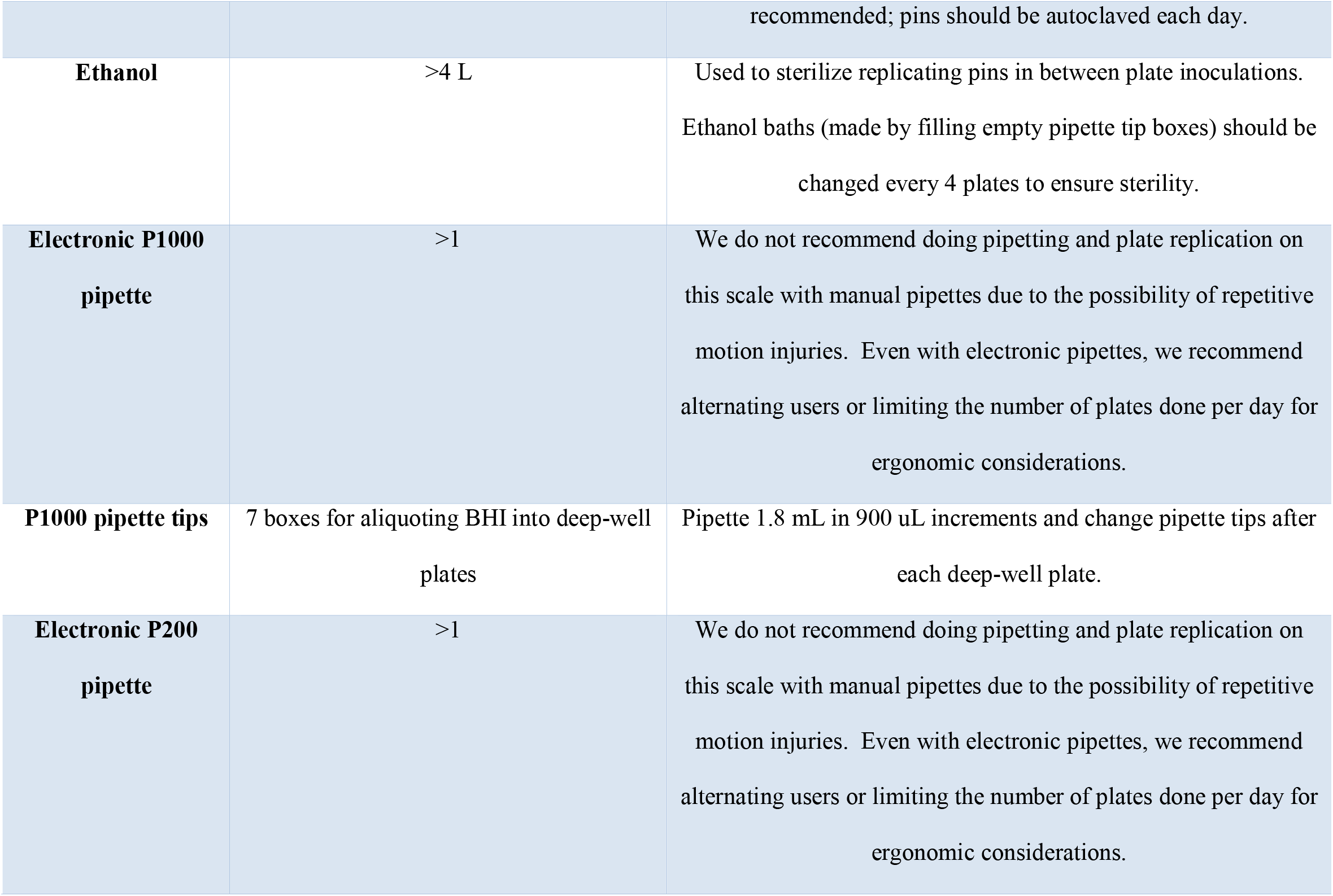
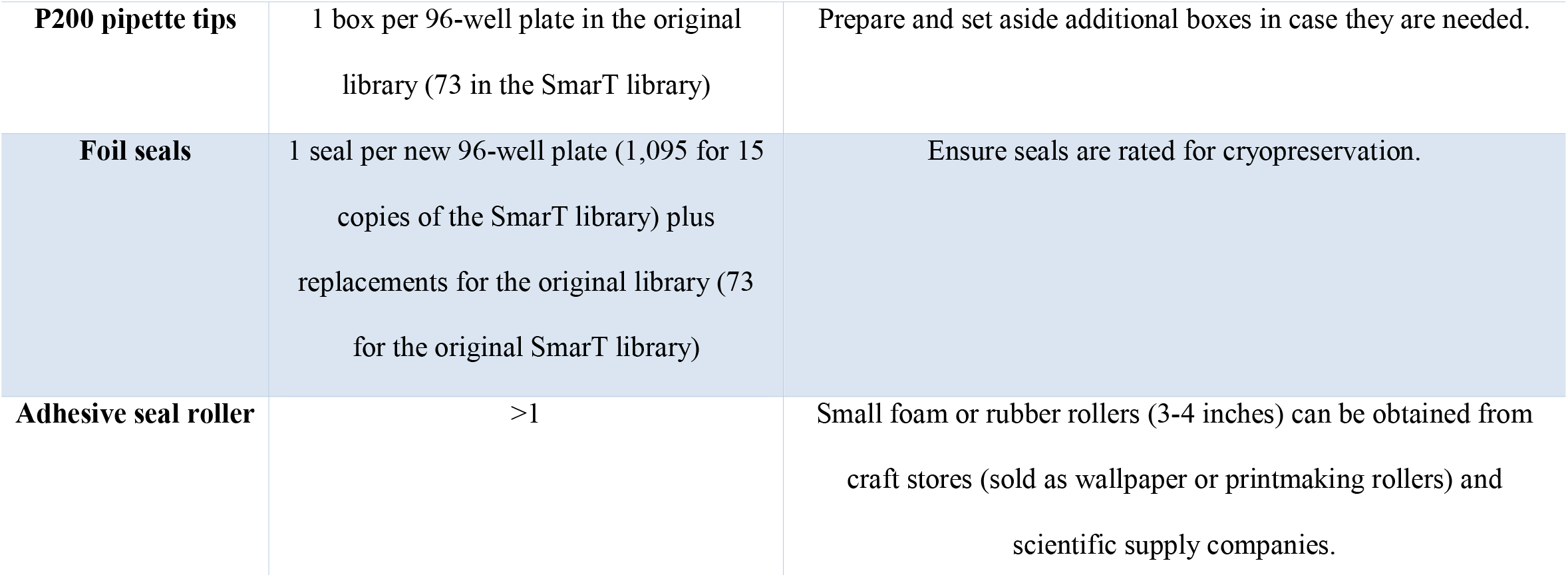
Required reagents, consumables, and equipment

**Table 2.** Timeline for generation of 15 copies of a 73-plate library

| **Task** | **Time** | **Notes** |
| --- | --- | --- |
| Ordering materials | variable | It is recommended to order ~10% extra of each reagent/supply |
| Generating plate labels | 4 hours | Ensure that cryogenic labels are used |
| Preparation of autoclaved reagents (BHI and glycerol) | ~8 hours | Prepared in 100 mL or 250 mL bottles to decrease chances of contaminating by repeated pipetting |
| Labeling and sorting 96-well plates | ~30 person hours for 1200 96-well plates |  |
| Distributing BHI into deep-well plates | ~35 person hours | Plus overnight incubation (done prior to inoculation to ensure no contamination) |
| Distributing 50% glycerol into 96-well plates | ~35 person hours |  |
| Inoculating | ~1 person hour per 4 96-well plates | Work in sets of 4 plates. Plates need ~10 minutes to thaw enough to use inoculating pins. This step requires overnight incubation. |
| Checking inoculated plates for blank wells and contamination | ~1 person hour per set of 16 plates (2 people x 0.5 hr) |  |
| Distributing cells from deep-well plate to new library copies & sealing with foil | ~1 person hour per deep-well plate (15 new 96-well plates) |  |
| Sorting and storing new library plates at -80 °C | ~2 person hours each day |  |

**Table 3.** Planned vs. actual timeline for replication of original plates in SmarT library

|  | Planned |  |  | Actual |  |  |
| --- | --- | --- | --- | --- | --- | --- |
|  | Transfer | Inoculate | Total complete | Transfer | Inoculate | Total complete |
| **Day 1** | - | 10 | - | - | 10 | - |
| **Day 2** | 8 | 16 | 8 | 8 | 16 | 8 |
| **Day 3** | 14 | 16 | 22 | 8 | 16 | 16 |
| **Day 4** | 14 | 16 | 36 | 10 | 22 | 26 |
| **Day 5** | 14 | - | 50 | 15 | - | 41 |
| **Day 6** | - | 16 | 50 | - | 16 | 41 |
| **Day 7** | 14 | Remainder | 64 | 16 | 17 | 57 |
| **Day 8** | Remainder | - | 73 | 14 | - | 70 |
| **Day 9** |  |  |  | - | 3 | 70 |
| **Day 10** |  |  |  | 3 | - | 73 |

**Figure 1.**
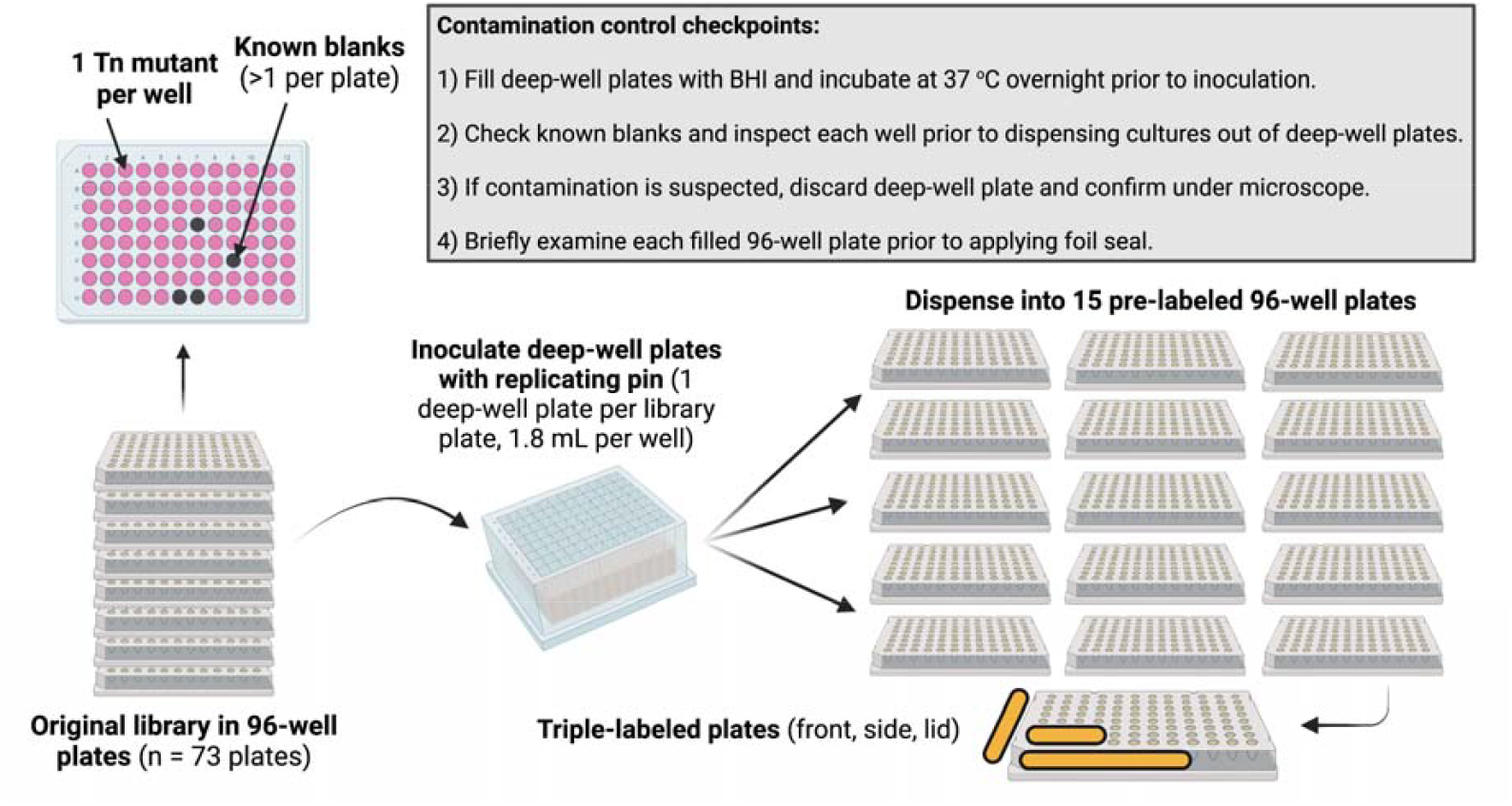
Overview of arrayed library replication and quality control checkpoints. SmarT Tn library plates were used to inoculate deep-well plates. Cultures were grown and examined for contamination, then dispensed into 15 individual pre-labeled, pre-loaded plates for new library sets. This image was created using BioRender.

Multiple quality control checkpoints were used to prevent contamination. The entire process was carried out in a Class 2 (A2) biological safety cabinet using BSL-2 practices. Deep-well plates were filled with BHI and incubated overnight at 37 °C prior to inoculation to ensure a lack of contamination. Replicating pins used for inoculation from freezer stock plates were sterilized in a series of ethanol baths in between plates. To ensure against dilution effects that can reduce the effectiveness of alcohol-based disinfection, the ethanol baths were completely replaced after inoculation of 4 deep-well plates. Replicating pins were also autoclaved each night.

The original SmarT library layout included a mapped series of known blank/empty wells in each plate (5). These were manually inspected in the deep-well plates, and any plates with contamination were discarded and reinoculated. Any deep-well plates where mutants did not grow as expected were also discarded and reinoculated. Because the deep-well plates do not fit in standard plate readers, optical density/absorbance was not measured. We recognize that as a limitation of this protocol, as that data could be important for identifying mutants with reduced overnight growth that is not detectable by eye. Importantly, we did not find any mutants in the entire 6,829-clone library that lost viability since the initial library construction.

New copies of library plates were pre-filled with glycerol and capped with sterile foil seals after addition of bacterial cultures. Plates were mixed by inversion after sealing (instead of pipetting) due to the number of samples. To ensure that this would not introduce cross-well contamination, we first empirically investigated the potential for cross-contamination. We filled a 96-well plate with glycerol, buffer, and bromophenol blue for visualization. This plate was covered with a foil seal, vortexed at 2500 rpm for 5 min, and incubated on a rotating shaker overnight. This agitation exceeded the brief mixing process done with Tn library plates containing cultures and glycerol. No dye leakage or damage to the foil seal was observed (**Figure 2**), suggesting that brief mixing of Tn library plates would not cause intra-plate contamination. This is consistent with our experience creating and handling the original arrayed Tn library.

**Figure 2.**
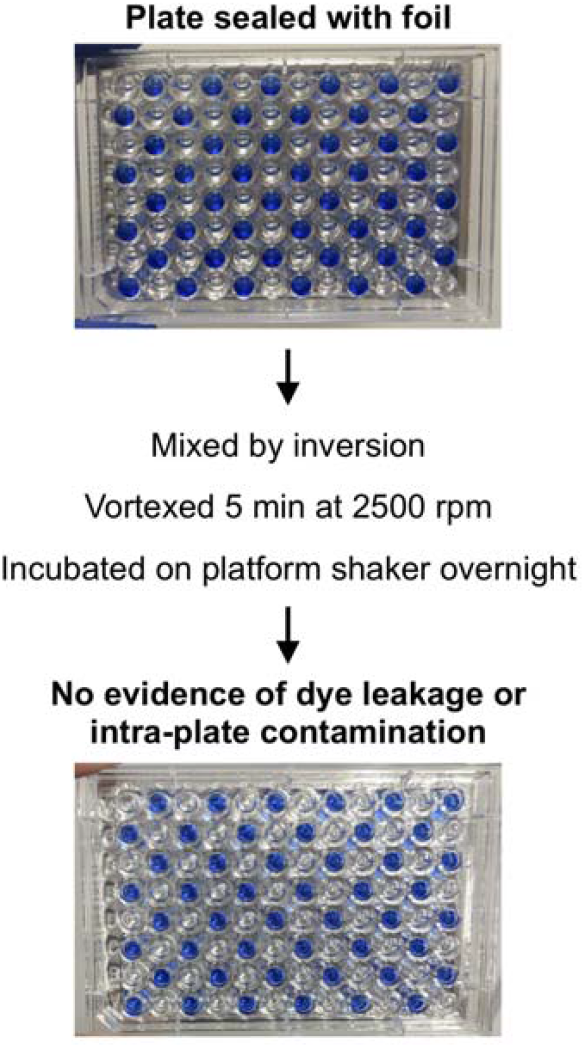
Sealing and mixing process does not introduce intra-plate contamination. A test plate with glycerol and bromophenol blue was sealed with foil, mixed by inversion, vortexed, and incubated on a shaking platform overnight to ensure that the inversion process used to mix glycerol and bacterial culture would not create contamination between wells.

### Pooling the SmarT Tn library from 96-well plates and Tn sequencing

The SmarT Tn libraries are available in arrayed format and as a pooled version, where all Tn mutants have been combined at equal amounts to facilitate TnSeq or similar genetic screens. Previously, the pooled versions were created by plating aliquots of each strain on BHI agar plates followed by scraping and combining all mutants (5). This was done to ensure that roughly the same number of mutants would be present in the pooled library regardless of *in vitro* growth defects in liquid medium and to screen each mutant stock for contamination. To determine whether we could pool liquid cultures of the SmarT library after growth in deep-well plates and still achieve a similar balance of mutants, we combined ∼200 uL remaining from each deep-well plate after distribution of the cultures into the new library plates. DNA was extracted from the pooled cultures, and Tn abundance was determined using Illumina sequencing and previously established protocols (5). These results were compared to samples extracted from the original pooled SmarT library in a previous experiment (17).

We first compared the number of Tn mutants identified from sequencing with the known number of mutants in the arrayed library (n=6,829). In the new pooled library, 250 mutants were missing (0 reads) in all extracted replicates (358, 336, and 347 mutants in the individual replicates) (**Supplementary Table 1**). 192 mutants had 0 reads in all previously sequenced samples (275, 259, 268, and 267 mutants in the individual replicates) (17). 166 mutants had 0 reads mapped in all replicates of the old and new pooled libraries. These mutants may be missing due to incomplete lysis of cells (perhaps due to physiological changes due to the disrupted gene), instability of the Tn insertion in the chromosome, or loss of DNA during preparation and sequencing. Because we did not find any mutants that did not grow in deep-well plates during library replication, we do not believe that these mutants are missing from the sequencing results due to a loss of viability. We next examined the similarity in relative abundance of mutants in the original pooled library compared to the new library pooled from liquid cultures (**Figure 3A**). In addition to a higher number of mutants with 0 reads, the new pooled library had a broader distribution of relative abundance frequencies (**Figure 3B**). Low abundance mutants in new pooled library (relative abundance 0 to 0.00001) have higher relative abundance in the original input library (**Figure 3C**). However, mutants absent from or with relatively low abundance in the original pooled library also had low abundance in the new pooled library (**Figure 3D**). Overall, we conclude that the original pooled library created by collecting cells grown on agar plates had a more even distribution of mutants than the new pooled library created by combining liquid cultures.

**Figure 3.**
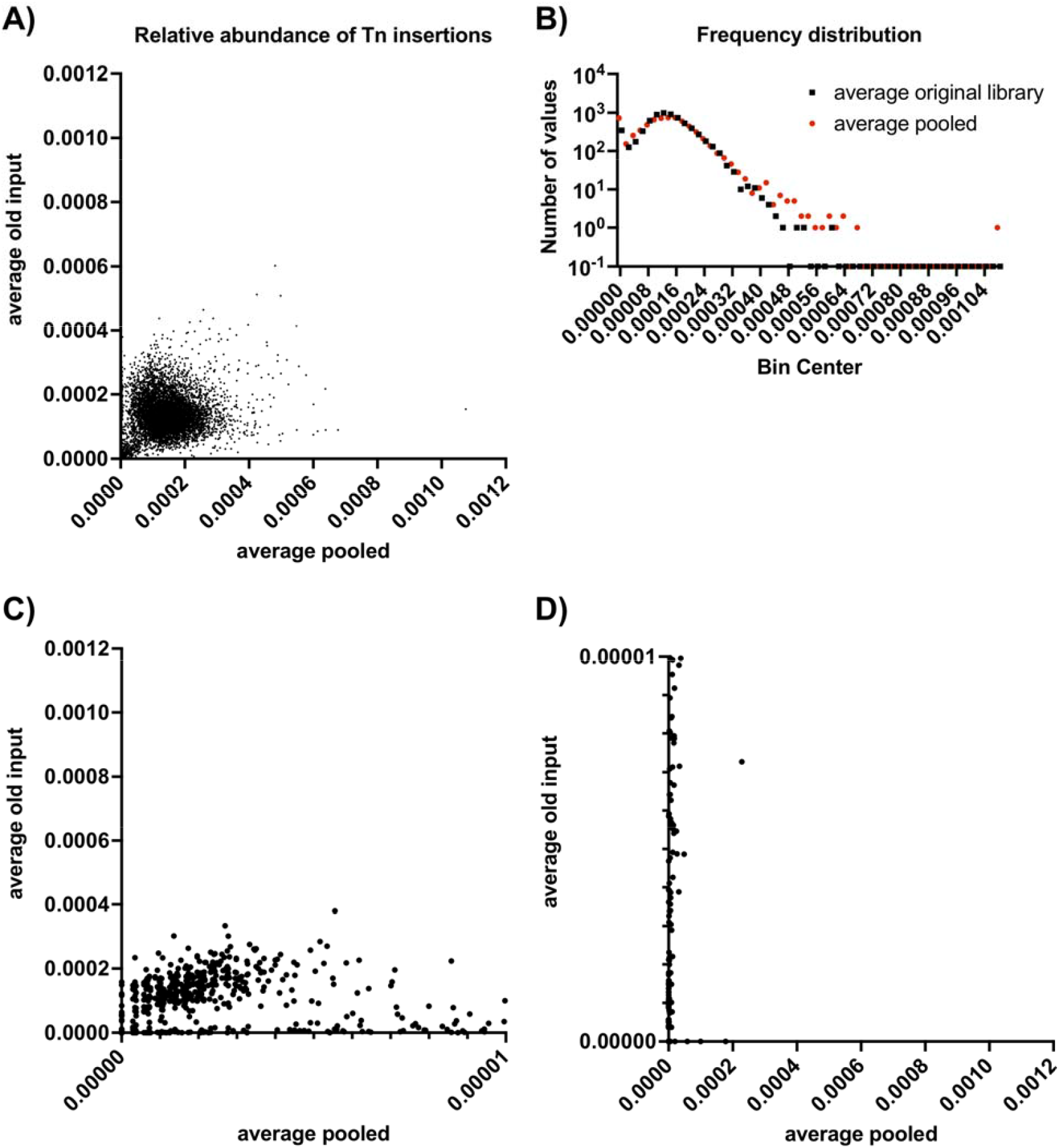
Comparison of SmarT library pooled from liquid cultures compared to original pooled library format. **A)** Relative abundance of Tn mutants in each library. **B)** Distribution of relative abundance. **C)** Low abundance mutants from new pooled library relative to abundance in original library. **D)** Low abundance mutants from original library relative to abundance in new pooled library.

## Discussion

Culture collections and arrayed mutant libraries are valuable biological resources that increase throughput, rigor, and reproducibility of experiments across an entire scientific field. To avoid redundancy and wasted resources, these collections and libraries should be broadly available to researchers. While core facilities or private companies may have resources to generate arrayed library copies using robotic arraying and liquid dispensing equipment, this remains beyond the reach of most academic research laboratories at many institutions.

Therefore, we sought to establish a protocol for manual replication of arrayed library collections that would increase accessibility to these biological resources while maintaining high quality control standards and preventing genetic drift due to repeated freeze/thaw cycles of arrayed culture collections.

Using this approach, we created 15 copies of a large arrayed *E. faecalis* OG1RF Tn library and have already distributed most of these library sets to other research groups. We also pooled Tn mutants grown during library replication and used TnSeq to compare mutant distribution from this pool to a previously generated pooled library in which individual mutants were scraped from agar plates. Although we found that a majority of Tn mutants (>96%) were present in our new pooled library, we observed greater representation of low-abundance mutants using the previous approach of scraping and pooling mutants from agar plates. Together, this methodology can guide the creation and distribution of arrayed mutant collections in a variety of microorganisms.

## Materials and Methods

### Bacterial strains and culture conditions

The 6,829 clone *E. faecalis* arrayed Tn library was previously generated and stored at -80 °C (5, 12). Brain Heart Infusion (BHI, BD Difco) was used for overnight growth. Prior to inoculation, plates were filled with BHI, incubated at 37 °C, manually inspected for contamination prior to inoculation. Tn mutants were inoculated using a metal replicating pin (Boekel Scientific) into 2 mL deep 96-well plates (Biotix) containing 1.8 mL BHI. Plates were grown without shaking overnight at 37 °C and manually inspected for contamination prior to distributing cultures to new library plates.

### Preparation of library plates

Sterile flat-bottom 96-well plates (Fisher Scientific) were labeled on the lid and two sides of the plate with printed cryovial labels (LabTags). Labels contained library copy number (1-15) and plate number (1-73). 100 uL of autoclaved 50% glycerol (VWR) was dispensed into each well using multichannel electronic pipettes.

### Generation of new library copies

From each deep-well plates, 100 uL of overnight cultures was distributed into 15 pre-labeled 96-well plates (pre-filled with 100 uL 50% glycerol) using multichannel electronic pipettes. Sterile AlumaSeal adhesive foil seals (Life Science Co) were applied with a small rubber roller (Speedball). Plates were sorted and stored at -80 °C.

### Preparation of pooled Tn samples for TnSeq and TnSeq analysis

200 uL was pooled from each well of each deep-well plate after distribution of mutants to new library plates. Samples were pooled in 50 mL Falcon tubes, pelleted at 6,000 rpm for 10 min in a Beckman Coulter Avanti JXN-30 floor centrifuge, and stored at -20 °C until further use. DNA was extracted using a Qiagen DNeasy Blood and Tissue Kit with a lysozyme pre-treatment step, as previously described (17), and submitted to the University of Minnesota Genomics Center. Libraries were prepared using a NEBNext Ultra II FS DNA Library Prep Kit for Illumina and a Nextera® XT Index Kit v2 Set A. Libraries were sequenced on a NextSeq P1 flow cell (150 PE) with ∼2.1 million reads per sample (∼300 reads/mutant). Mutant abundance was quantified using custom scripts as previously described (5, 17).

## Acknowledgements

This work was supported by a grant from Boehringer Ingelheim Fonds to AL, American Heart Association fellowship 907592 to DB, National Institutes of Health grant K99AI151080 to JLEW, and National Institutes of Health grant R01AI122742 to GMD. The authors thank Cristel Archambaud and Pascale Serror for supporting the participation of AL and Jose Lemos for supporting the participation of DB. The authors are also grateful to Pascale Serror and Jose Lemos for helpful comments on the manuscript.

