## Supplementary File 1 for "Optimized replication of arrayed bacterial mutant libraries increase access to biological resources"

**Detailed protocol**

1. Label deep-well plates using a marker or printed labels.
2. Using printed labels, label 96-well plates that will be used for new library copies. Affix labels to the lid and side of each plate so that at least one label is present on the base when lid is removed.
3. Prepare and autoclave 50% glycerol for the 96-well plates (calculate required volume based on 100 uL per well).
4. Using an electronic pipettor, aliquot 100 uL autoclaved 50% glycerol into each well.
   a. These can be prepared in advance and stored at room temperature.
   b. If possible, do this step in a biosafety cabinet to prevent contamination.
5. Autoclave required volumes of BHI (or other growth medium) and 50% glycerol. Aliquot medium into deep-well (≥2 mL total capacity) 96-well plates. Incubate at 37 ^o^C overnight. Examine plates the next day to make sure there is no contamination.
   a. We recommend using 100 uL per library copy plus 20% extra volume in case of evaporation or pipetting error. For 15 library copies, we added 1.8 mL to each deep-well plate.
   b. If possible, do this step in a biosafety cabinet to prevent contamination.
6. If replicating an existing Tn library that is stored at -80 ^o^C, remove frozen Tn library plates from the freezer in batches of 4 plates and transfer to a biosafety cabinet. Remove foil seal covering plates. Remove the foil seal while the plate is still frozen to reduce chances of cross-contamination between wells. Thaw the frozen plates enough to use a replicating pin. Avoid bringing plates all the way to room temperature. In our experience, this takes 8-12 minutes.
7. Using a 96-well replicating pin, inoculate the deep well plates containing BHI. Sterilize the replicating pin in between plates using a series of ethanol baths.
   a. You should be able to push the replicating pin to the bottom of the plate to ensure the ends of all pins are submerged. If plates have not thawed enough, wait 1-2 minutes and try again. Using excess force with the replicating pin can damage the Tn library plate, resulting in cross contamination.
   b. Replace the ethanol baths after every set of 4 plates.
8. Incubate the inoculated deep-well plates at 37 ^o^C overnight.
9. After overnight growth, check the deep-well plates to confirm growth of all mutants and that the known blank wells are not contaminated. If any mutants did not grow or any wells are contaminated, discard the entire plate and inoculate a fresh plate.
10. Using a multichannel electronic pipettor, transfer 100 uL out of the deep-well plates into the corresponding 96-well plates containing glycerol.
11. Cover each plate with a foil seal. Use the rubber roller to form a tight seal around the edges and over the top of the plate.
12. Sort into bags or storage containers and store Tn library copies at -80 ^o^C.
